# Quantitative evaluation of enrichment protocols on rhesus macaques welfare in laboratory environment

**DOI:** 10.64898/2026.06.10.730846

**Authors:** Valentine Morel-Latour, Océane Monnier, Elya Ray, Alyssia Fort, Paul Roux, Manon Dirheimer, David Thura, Fadila Hadj-Bouziane, Charles R. E. Wilson, Emmanuel Procyk, Jérôme Sallet

**Affiliations:** Université Lyon 1, INSERM, SBRI, U1208, Lyon, France; Inserm, Délégation Régionale Auvergne Rhône-Alpes, 69500 Bron, France; Université Lyon 1, Lyon, France; Centre de Recherche en Neurosciences de Lyon, Inserm U1028, CNRS UMR5292, Lyon, France; VetAgroSup, Marcy l’Etoile, France; Réseau médical & scientifique de l’Institut de Neuromodulation, INM, GHU Paris Psychiatrie et Neurosciences; Wellcome Centre for Integrative Neuroimaging, Department of Experimental Psychology, University of Oxford, Oxford, United Kingdom

**Keywords:** Rhesus macaques, Housing, Enrichment, Welfare, Behaviour

## Abstract

Non-human primates are crucial animal models in biomedical research including neurosciences. Legal frameworks ensure that participation of monkeys in biomedical research follows an ethical assessment, driven by specific guidelines for ensuring animal well-being. Those guidelines promote continued refinement of practices to improve animal wellbeing, yet quantitative data to inform such refinements is limited. In this context, we investigated the impact of modifications to the housing that created either navigational complexity or visual access to neighbouring animals on the behaviour of rhesus macaques. Then we quantified the value of different enrichment programmes based on manipulable objects. We showed that making navigation within a modular housing more complex, or adding transparent separations between housing units is associated with significant reduction of severe aggression, stereotypical behaviours and a significant increase of appeasing behaviours. We also observed a positive effect of manipulable objects with an increase in the expression of appeasing behaviours and a decrease in abnormal or stereotypical behaviours. Finally, our study revealed significant inter-individual variability that was not explained by sex, age, social network size or social status. Overall, our results point to the importance and benefits of environmental changes and enrichment protocols, and of the critical need for personalised enrichment programmes.

## Introduction

Non-human primates are crucial animal models in biomedical research (2023, Washington (DC), National Academies Press (US);,1–4). They contribute to the advancements of fundamental and clinical research in many fields including immunology, reproductive health, and neuroscience, when scientific questions target primate specific mechanisms and no replacement is available. In neuroscience, non-human primate models are essential for understanding cerebral function and developing new therapies, as for example deep brain stimulation or brain computer interface-based approaches (1,5). Simultaneously, animal experimentation also raises important ethical questions for researchers and civil society alike. Legal frameworks like the European Directive 2010/63/UE (revision of the directive n°86/609/CEE), ensure that the participation of monkeys in biomedical research follows an ethical assessment, and is performed under specific housing conditions, with enrichment and veterinary monitoring adapted to the species and the individual (for review see 6).

To promote the welfare of animals involved in scientific procedures, environmental enrichments, that are conceived as positive changes for the animal from the human’s point of view, are used (7). Those changes could reflect the structural organisation of the enclosures with the addition of swings or perches, the provision of a foraging substrate and the provision of manipulable objects (8,9). Despite this diversity, all enrichment serves one purpose. They must enable animals to make better, more diverse or richer use of their environment, or at least provide them with greater control over it through choice, enabling them to express a varied repertoire of natural behaviours. The impact of an enrichment can be measured through an animal’s interest for it, by measuring the time it spends interacting with that enrichment (10,11), its effect on the diversity of behaviours expressed by the animals (12,13) or its effects on the animal’s physiology (14,15). It is also possible to measure indirect effects such as a reduction of abnormal behaviours (e.g. pacing), a modulation of the time budget or a reduction in aggression within a social group (16–18).

Aggressive and abnormal behaviours (e.g. pacing) are arguably the most common potential issues to be addressed with captive animals. Enrichment programmes are used to prevent or to reduce those behaviours. Considering the invasiveness of research, providing more space, or more toys are too simplistic approaches to questions of aggressivity and abnormal behaviours in captive primates (19). To best ensure development and implementation of solutions for improving animal welfare, one would need good quality quantitative data on the benefits of enrichment interventions for captive monkeys. Furthermore, solutions should be, if possible, cheap and easy to use for maximizing their implementation as widely as possible. Finally, chosen solutions should maximize the welfare of each individual. Thus, the inter-individual variability in responses to enrichment requires to include individual differences in assessing enrichment programmes (20). Developing an adapted enrichment programme is crucial, especially for animals that for reasons of behavioural incompatibility or on veterinary advice have to be separated from their social groups. In this project, we tested the impact of two simple structural modifications of the primate housing on animal welfare. We also investigated the benefit of a cumulative provision of manipulable objects, and the inter-individual variability with regards to preferences for those objects.

Overall, our behavioural results showed several positive impacts of our modifications to the primate housing on animals’ wellbeing. First of all, we showed that by adding more navigational paths in animal enclosures, we were able to change dynamics in the social groups. The use of transparent separation in modular housing reduced pacing, and promoted pseudo-social behaviour in rhesus macaques as previously observed in cynomolgus macaques (*macaca fascicularis*) (15). Finally, we showed that attention should be paid to proposing enrichments adapted to each individual when designing an enrichment programme.

## Material and methods

### 1. Subjects

A total of 20 rhesus macaques from 3 research colonies were included in 4 studies aiming to quantify the impact of several manipulations of the primate housing environment on primate welfare. This project was discussed with the Animal Welfare Body of the laboratories in which animals were housed. The data presented in this article were collected in the context of ethically approved protocols by French Animal Experimentation Ethics Committee #42 (CELYNE) that followed guidelines of the European Community on animal care (European Community Council Directive 2010/63EU).

Animals were housed in modular compounds, within which animals were assigned to distinct social groups of two to three animals. Note that 7 animals lived in individual pens due to social incompatibility issues (see Table 1) but those animals were housed in rooms with other monkeys.

**Table 1:** Summary of information about all monkeys tested and the studies they were implicated in.

| ID | Sex | Age (at the time of data collection) | Study | Social Group Size | Number of observation sessions | Relative Social Status (% dominant behaviour) | Colony ID |
| --- | --- | --- | --- | --- | --- | --- | --- |
| Bo | M | 7 | 1 ; 3 ; 4 | 3 | 24 | 62 | 1 |
| Fa | M | 7 | 1 ; 3 ; 4 | 3 | 24 | -55 | 1 |
| Dj | M | 6 | 1 ; 3 ; 4 | 3 | 24 | 2.7 | 1 |
| Dai | F | 14 | 2 ; 3 ; 4 | 1 | 24 | NA | 1 |
| Ra | M | 11 | 2 ; 3 ; 4 | 1 | 23 | NA | 1 |
| Lo | M | 14 | 1 ; 4 | 2 | 21 | 26.6 | 1 |
| Na | F | 14 | 1 ; 4 | 2 | 21 | -26.6 | 1 |
| Dal | M | 18 | 1 ; 4 | 2 | 33 | 62.5 | 2 |
| Pi | F | 18 | 1 ; 4 | 2 | 33 | -62.5 | 2 |
| Is | F | 12 | 4 | 2 | 14 | -60.8 | 2 |
| Gu | F | 14 | 4 | 2 | 14 | 60.8 | 2 |
| Qu | M | 13 | 2 ; 4 | 1 | 14 | NA | 2 |
| Jf | M | 11 | 4 | 2 | 20 | 68.2 | 2 |
| Js | F | 11 | 4 | 2 | 20 | -68.2 | 2 |
| Li | F | 21 | 4 | 1 | 18 | NA | 3 |
| Ce | F | 25 | 4 | 1 | 19 | NA | 3 |
| Am | F | 13 | 4 | 2 | 20 | 40 | 3 |
| Eo | F | 13 | 4 | 2 | 20 | -40 | 3 |
| Di | M | 6 | 4 | 1 | 14 | NA | 3 |
| Bi | M | 6 | 4 | 1 | 10 | NA | 3 |

Seven rhesus macaques (macaca mulatta) (five males, two females; 12,0 ± 5,0 years old; Table 1) participated in the first study (Study one - Modification of enclosures for socially-housed monkeys).

Three rhesus macaques (macaca mulatta) (two males, one female; 12,7 ± 1,5 years old; Table 1) were subjects for the second study (Study two - Modification of enclosures for single-housed monkeys).

Five rhesus macaques (macaca mulatta) (four males, one female; 9,0 ± 3,3 years old; Table 1) were subjects for the third study (Study three - Impact of enrichments on socially and single housed monkeys’ time budget).

Finally, twenty rhesus macaques (*macaca mulatta*) (ten males, ten females; 12,2 ± 4,8 years old; Table 1) were involved in the last study (Study four - Object relevance for individualised enrichment programmes).

No animals were excluded from any of the four studies. Sample size was constrained by the number of animals and the housing conditions in the research facility at the time of the project was conducted. Due to the limited number of subjects, one could consider Study 1 to 3 as pilot studies.

The relative dominance status index was closely based on the method used in previous studies (21–23). The directions of agonistic behaviours (aggressive/dominant or submissive) of 13 socially-housed animals were recorded during a series of 5-minute observations (each preceded by a 5-minute habituation period; Table 1). Behaviours recorded include the following social behaviours: chasing, escaping, aggressive and non-specific social behaviours (i.e. resting together). Because it has been suggested that grooming and mounting have no necessary relationships with dominance these behaviours were classified as non-specific social behaviours (24). In addition, non-social behaviours were also recorded during these sessions. They included resting, foraging, playing, scratching and self-grooming. Non-social behavioural observations were also collected for single-housed animals. The number of observation sessions per social group ranged from 14 to 33 and from 10 to 24 for socially-housed animals and single-housed animals respectively. Observations were conducted by six independent observers (VML, OM, ER, JS, OJ, MV).

Those observations were used also in Study 1 to evaluate a putative change in behaviour associated with changes of the primate housing.

### 2. Experimental protocols

All the videos collected were analysed with the software BORIS (Behaviour Observation Research Interactive Software) (25). All data were analysed using R (R Core Team (2021). R: A language and environment for statistical computing. R Foundation for Statistical Computing, Vienna, Austria, https://www.R-project.org/).

### 2.1 Study one - Modification of enclosures for socially-housed monkeys

#### 2.1.1 Behavioural observations

This study focused on socially-housed monkeys. European Directive 2010/63/EU provides guidelines on minimal amount of space per enclosure (3.6m3), and per animal (1.8m3). Despite being provided with more than the recommended standards, social tensions could occur and lead to severe injuries. Here, we tested whether offering not more space, but more options to escape from other monkeys could impact on monkey welfare. More specifically, we studied the impact of complexifying navigational pathways in primate housing by removing separation panels on different levels of the cages, allowing for the creation of circuits (Fig. 2A). We hypothesized that offering multiple options to animals for navigating through their housing could help to reduce social tensions in the group when they arise.

For this study, we analysed behavioural data collected from ethograms in a social group of 3 monkeys (G1: Bo, Fa, Dj), and a pair of animals (G2: Dal, Pi). We analysed three categories of behaviours. The first one named “appeasing non-social behaviours” included foraging, playing with/using an object or a toy, and self-scratching/grooming behaviours. The other two categories were related to social behaviours, either behaviours indicative of social tension in the group, or of social appeasement. We considered aggression, attack, and escape behaviours in the “aggressive social behaviours” category and grooming or resting together in the “appeasing social behaviours” category. Each animal was its own control.

The occurrence of each behaviour was measured during 5-minute observation periods during which circuits were ON (all panels removed, offering two paths to navigate between adjacent enclosures) or OFF (half of the panels removed, offering only one path to go from one enclosure to the next). We collected data from 8 and 10 observation sessions per social group prior to the introduction of circuits in G1 and G2 respectively (Circuit OFF). We then collected 12 and 11 observation sessions after the introduction of circuits in these two groups (Circuit ON). Finally, we performed an assessment of the occurrence of injuries prior versus after the implementation of circuits. We added data from a third social group (G3: Lo, Na) for this qualitative evaluation.

#### 2.1.2 Statistical analysis

Analysis focused on the effect of the condition (circuit ON or circuit OFF) on three categories of behaviours measured. For “appeasing non-social behaviour”, we fitted a generalised linear mixed model (GLMM) using a negative binomial distribution to account for unbalanced sample size (58 values for Circuit ON and 44 for Circuit OFF), repeated measures (multiple sessions per monkey, var = 0.92), not normally distributed residuals and overdispersion (mean = 2.96; var = 6.16; ratio > 2). The dependent variable was appeasing non-social behaviour, with condition included as a fixed effect and monkey identity as a random effect.

For “appeasing social behaviour”, we fitted a generalized linear model with Poisson distribution because residuals weren’t normally distributed, there was no overdispersion (mean = 0.19; var = 0.21; Mean ≈ Var), there were no excessive zeros. The dependent variable was appeasing social behaviour, with condition included as a fixed effect.

Finally, for “aggressive social behaviour”, we fitted a generalised linear mixed model (GLMM) using a negative binomial distribution to account for unbalanced sample size, repeated measures (var = 0.19), not normally distributed residuals and overdispersion (mean = 0.55; var = 1.60; ratio > 2). The dependent variable was aggressive social behaviour, with condition included as a fixed effect and monkey identity as a random effect.

### 2.2 Study two - Modification of enclosures for single-housed monkeys

#### 2.2.1 Behavioural observations

This study was specifically designed for singled-housed monkeys. We tested the impact of a structural enrichment on monkeys’ behaviours by replacing the opaque panel between a single-housed monkey and their neighbour with a transparent panel. This work was inspired by the study conducted by Disarbois and colleagues on cynomolgus monkeys (15).

We measured in three monkeys the duration of stereotypical pacing behaviour during 10-minute observation sessions with three different conditions. In typical setting conditions, the opaque panel was set permanently (condition *Opaque*). Then, we introduced the transparent panel over a 4-week period. In the first 2.5 weeks, the transparent panel was introduced 2 to 3 hours per day (condition *Habituation*), followed by 1.5 weeks during which the transparent panel was put in the morning and removed at the end of the day. After the habituation phase, the transparent panel was set permanently (condition *Transparent*).

In each condition, behaviours were recorded using a GoPro for 10 minutes, two to three times per day during a week. Altogether, we collected 14 observations for monkeys Dai and Ra, and 12 observations for monkey Qu in the condition *Opaque*. For these animals we collected 12, 12 and 11 observation sessions in the condition *Habituation* and 13 observation sessions in the condition *Transparent* for the monkeys Dai and Ra. No observation session was conducted for Qu in the condition *Transparent* as Qu was rehoused in a different room during the project. Each animal was its own control.

#### 2.2.2 Statistical analysis

Analysis focused on the effect of the condition (condition *Opaque*; condition *Habituation*; condition *Transparent*) on the time spent pacing. We fitted a linear mixed effects model (LMM) because: sample sizes were unbalanced (40 values for condition *Opaque*, 35 values for condition *Habituation* and 26 values for condition *Transparent*), repeated measures were made and residuals were normally distributed. We then conducted pairwise comparisons with Bonferroni correction. The dependent variable was pacing duration, with condition included as a fixed effect and monkey identity as a random effect.

### 2.3 Study three - Impact of enrichments on socially and single housed monkeys’ time budget

#### 2.3.1 Behavioural observations

This study evaluated the impact of a cumulative enrichment programmes on the time budget of 5 monkeys with 3 different conditions: the first one without giving additional objects to the monkeys (condition *Without*), a second one with the distribution of known objects (condition *Known*), a third one with the distribution of only new objects that monkeys never saw before (condition *New*). Data were collected on 3 socially-housed monkeys and 2 single-housed monkeys.

On day 1, monkeys were fed in the morning, then they received two enrichments at 3pm (PM1). Their behaviours were recorded using one or two GoPros for 10 minutes. Two hours later (PM2), a second recording was made before feeding and foraging mix was given to the monkeys. On days 2 to 5, one recording was made in the morning (AM) before feeding and foraging mix, the second recording was at 3pm (PM1) at the time of the distribution of an enrichment, and the third recording was two hours later (PM2) before afternoon feeding and foraging mix distribution. All objects distributed are described in figure 1B. For the condition *Without*, recording hours were matched to the other two conditions (Fig. 1A).

**Figure 1:**
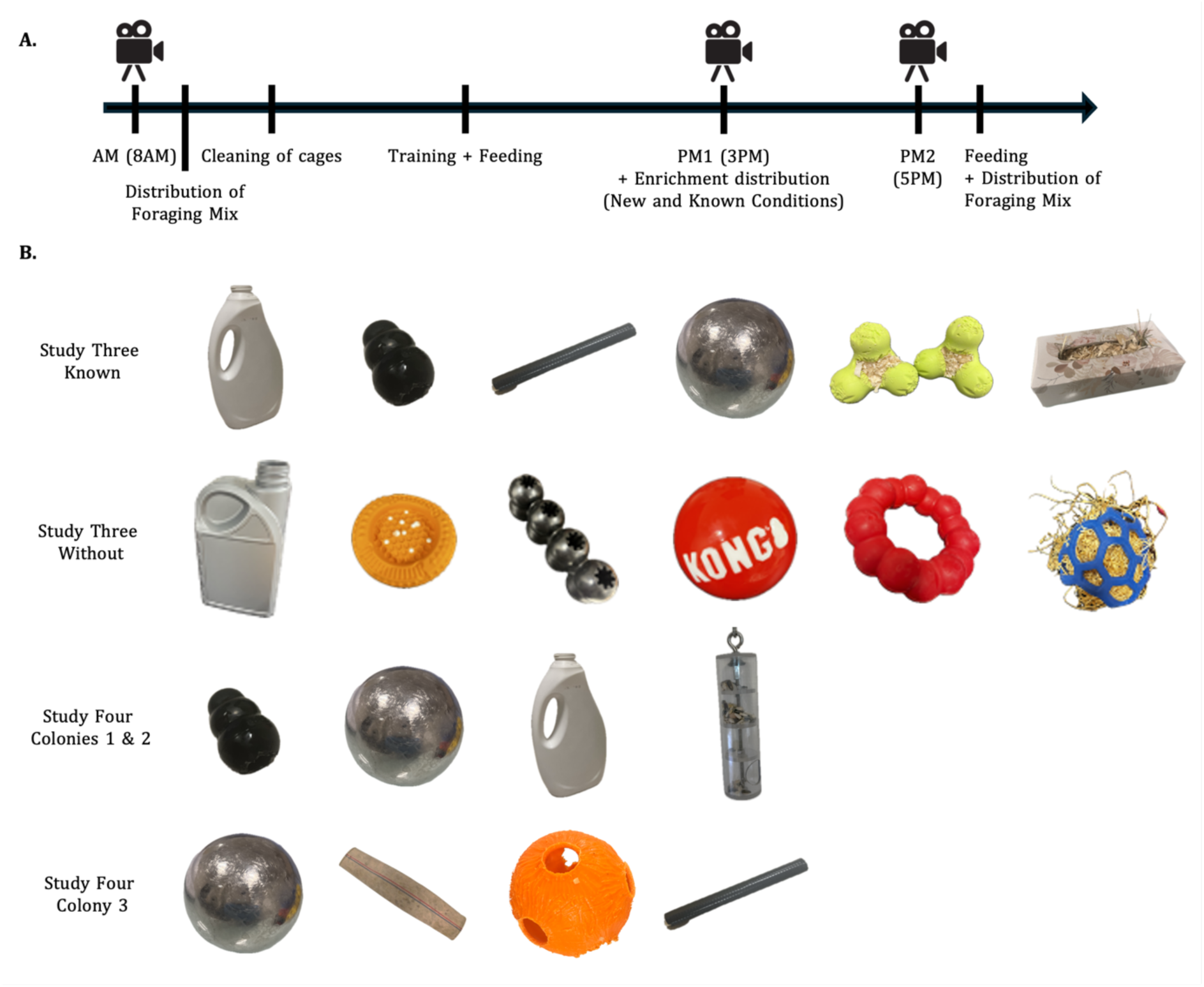
A. Timeline of study three describing the different steps of the days during which data were collected. Recordings were made at AM (8AM), PM1 (3PM) and PM2 (5PM). B. Pictures of all objects used in study three and four. For study three - Impact of enrichments on socially and single housed monkeys’ time budget, we used 6 objects per week and per monkey for the known objects and the new object conditions. For study four - object relevance for individualised enrichment programmes, we used seven different objects in total.

**Figure 2:**
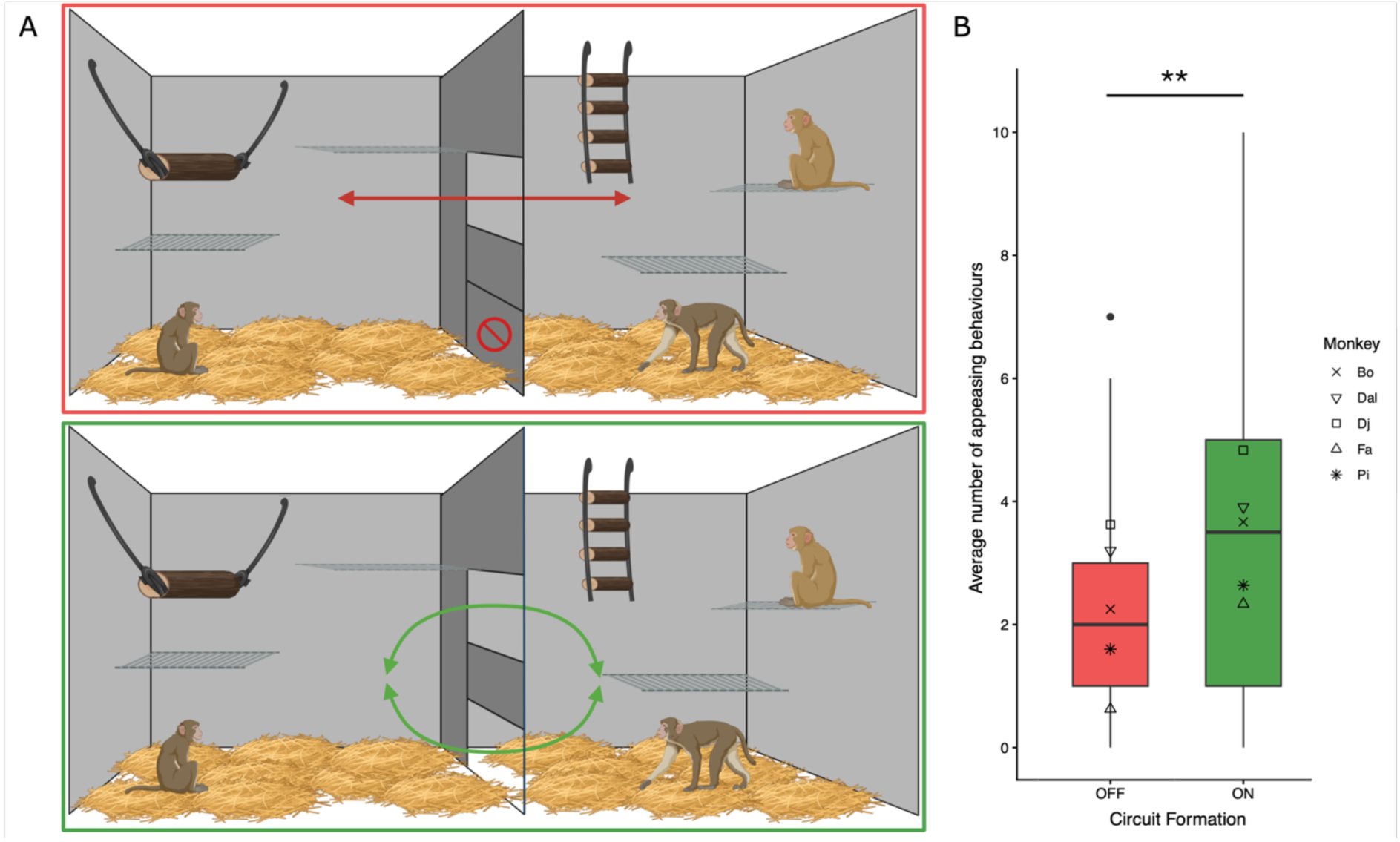
A. Schematic representation cages for socially-housed monkeys, without creation of circuits (top row) and with the creation of circuits (bottom row); figure created using biorender.com B. Number of appeasing behaviours across 5-minutes observation with and without the formation of circuits, averaged across five monkeys, shows a significant difference (p-value = 0.0066).

To study the general impact of these three conditions, we used the following ethogram composed of ten categories of behaviours: 1. affiliative and neutral social behaviour (presentation; mounting; grooming; affiliative; hug; resting together) 2. aggressive social behaviour (push away; avoid; attack; escape) 3. resting (resting alone; attention to outside: attention to other) 4. self grooming/scratching 5. feeding behaviours (eat; drink; forage) 6. moving 7. using object (1 to 7) 8. interaction with cage elements (structural enrichment; cage) 9. abnormal behaviour (pacing; self harm; pull hair; other) 10. other (sleep; yawn; call; pee/poop; shake cage; not visible). We collected in the same animals (Bo, Fa, Dj, Dai and Ra), 14 observations per monkey in the New and Without conditions and 15 observations per monkey in the condition *Known*. Each animal was its own control.

#### 2.3.2 Statistical analysis

For this study, we conducted our analysis first at the group level to see if there was a general effect of the condition on monkeys’ time budget. As interindividual variability is strongly reported here, we also conducted a subject-by-subject analysis, testing how the three conditions impacted on ten behaviours for each individual.

##### 2.3.2.1 Group Level - Permanova

Some behaviours are never or rarely observed in certain individuals, our dataset contains a high proportion of zeros (843 observations < 0.5 out of 2150). We applied a Hellinger transformation prior to the computation of Euclidean distances in order to reduce distortions arising from absent or rare behaviours. We then tested the dispersion homogeneity (betadisper). The test was significant (p-value = 0.002087), indicating that within-group variances were not homogeneous across conditions.

We conducted a Principal Coordinates Analysis (PCoA ordination) revealing that the three conditions (*New*, *Know*, *Without*) are not visually distinguishable. The points of the three groups blend together in the multivariate space, suggesting that behavioural profiles are broadly similar across conditions. The structure of the data was primarily driven by individual identity rather than condition.

We performed a permutational multivariate analysis of variance (PERMANOVA) to assess the effects of condition and time of the day on the duration of each behaviour after Hellinger transformation, using Euclidean distances and where permutations were restricted within individuals to account for the repeated-measures structure of the data. Marginal effects were tested using 9 999 permutations. We tested for marginal (independent) effect and sequential effects

As the values were almost identical (Condition by “terms”: F = 9.88, R² = 0.076 and by “margin”: F = 10.01, R² = 0.077; Time by “terms”: F = 14.84, R² = 0.114 and by “margin”: F = 14.85, R² = 0.114), it confirms that Condition and Time were not correlated in the experimental design. Permanova results need to be interpreted carefully because the dispersion homogeneity test was significant.

For pairwise post hoc comparison, we conducted the same permanova as the previous model but pair by pair and did Bonferroni correction.

##### 2.3.2.2 Subject-by-subject - Five models tested

For our analysis at the subject level, we then designed five different models (described below) that we tested on each behaviour of each subject and compared all fitted models using quality checks (DHARMa diagnosis) such as uniformity test, dispersion test and zero inflation test. We also calculated quality scores AIC and BIC. Then, the model with the highest validation score (DHARMa) and the better AIC was selected for the current behaviour (See Supplementary Table 1 for details on the model used per behaviour).

Models tested:

1. Beta with interaction (beta_interaction) → The variable duration of the tested behaviour (expressed as a percentage) was first converted to a proportion. To meet the assumptions of beta regression (i.e., values strictly bounded between 0 and 1), proportions were adjusted using the transformation (y * (n − 1) + 0.5) / n, where n is the sample size. Then, we fitted a generalized linear mixed-effects model (GLMM) with a beta distribution to model the transformed duration as a function of condition, time of the day, and their interaction, including the date as a random effect.
2. Beta without interaction (beta_additif) → We fitted a generalized linear mixed-effects model with a beta distribution to model the same transformed duration as a function of condition and time of the day, including the date as a random effect.
3. Zero-Inflated Beta with interaction (zi_beta_interaction) → We fitted a zero-inflated generalized linear mixed-effects model with a beta distribution to model the same transformed duration as a function of condition, time of the day, and their interaction, including the date as a random effect. A zero-inflation component (intercept-only) was included to account for excess zeros in the data.
4. Zero-Inflated Beta without interaction (zi_beta_additif) → We fitted a zero-inflated generalized linear mixed-effects model with a beta distribution to model the same transformed duration as a function of condition and time of the day, including the date as a random effect.
5. Gaussien (transformation sqrt) (gaussian_sqrt) → The variable duration of the tested behaviour was square-root transformed to improve normality and homoscedasticity of residuals. Then, we fitted a linear mixed-effects model (LMM) with Gaussian errors to model the transformed duration as a function of condition, time of the day, and their interaction, including the date as a random effect.

We conducted a type II anova on the best model selected followed by posthoc comparisons and Bonferroni correction.

### 2.4 Study four - Object relevance for individualised enrichment programmes

#### 2.4.1 Behavioural observations

To assess individual preferences, we gave an object to a monkey and recorded videos of 10 minutes using one or two GoPros. We then measured how long the object was used for. Twenty monkeys were tested, and 3 to 4 objects were presented to each individual: kong, silver ball, bottle, puzzle for colonies 1 and 3 and silver ball, fire hose, noise ball, tube for colony 2 (Fig. 1B). There was no control group in study four.

#### 2.4.2 Statistical analysis

Each subject was tested several times (repeated measures), observations are not independent. Values were confined between 0 and 100, data were close to the limits, and the distribution was skewed to the right (median = 13.965 < mean = 30.280). Data contained too many zeros and 100s strict (13 observations at zeros out of 78 and 2 observations at 100 out of 78) that could not be ignored nor transformed. We normalised data between 0 and 1 and chose a generalized linear mixed model with Beta Zero-One-Inflated (ZOIB) distribution containing all the variables we wanted to test (age, sex, social group size and social status) and controlling for interindividual variability, using monkey identity as a random factor. We also controlled for the potential effect of the nature of the object on the time of use but the model was not converging properly and the variance of the random effect object was close to zero so we removed it from the model.

As seven out of twenty monkeys were singly housed, we weren’t able to measure their social status. So, we tested 2 models: one including the social status of thirteen subjects (Model a) and one without the social status including all twenty monkeys (Model b, see below).

Models: a) We fitted a generalized linear mixed-effects model (GLMM) with an ordered beta distribution to model the time of use normalised as a function of age, sex, social group size, and social status, including monkey identity as a random effect.

b) We fitted a generalized linear mixed-effects model (GLMM) with an ordered beta distribution to model the time of use normalised as a function of age, sex, and social group size, including monkey identity as a random effect.

## Results

## 1. Structural modifications of the environment

### 1.1 Study one - Modification of enclosures for socially-housed monkeys

Monkeys are housed in modular enclosures. The use of moveable panels allows the organization of space to adjust for different group sizes, and also to give access to specific parts of the enclosure to technicians for cleaning procedures, or to animals for training purposes. When not needed, panels are removed and macaques can move across different cages.

Here, we tested if removing panels in such a way that it multiplies the paths available to monkeys to navigate across their enclosures could impact on aggressivity level and rate of injuries in groups of rhesus macaques. We referred to those paths as circuits (Fig. 2A). We hypothesized that it could provide animals with better coping options in case of social tension. To quantify the putative benefits of those circuits, we recorded behaviours of five monkeys in 102 observation sessions of 5 minutes, based on a protocol used in previous studies on social behaviours in macaques (21–23). 44 sessions were collected prior to the introductions of circuits. 58 sessions were collected following the introduction of circuits. We focused our analysis on three categories of behaviour. Behaviour indicative of social tension in the group (aggression, attack, escape), social appeasement (grooming, resting together), and non-social appeasing behaviours (playing or using an object/toy, foraging and self-grooming/scratching). We observed no difference in the occurrence of aggressive behaviours (p-value = 0.4639), or of socially appeasing behaviours (p-value = 0.0865). However, macaques engaged in significantly more appeasing behaviours when we offer them different navigation routes compared to when we don’t (p-value = 0.0066; Fig. 2).

We implemented circuits in G1 enclosures approximately 6 months after animals’ arrival in the facility. Since the implementation of circuits, more than 3 years prior to this study, 3 veterinary interventions were required on the most submissive animal in the 3-male social group (G1). While 4 interventions were required in the G2 pair of animals over a 2-year period prior to the implementation of circuits, no interventions were necessary for the last 1.5 years since the permanent implementation of circuits in their enclosure. Finally, in the G3 pair of animals, we compared the clinical history for 1.5 years prior to the implementation of circuits versus a more than 2-year period after their implementation. While several injuries caused by the dominant animal occurred when no circuit was available, no injury requiring a veterinary intervention occurred after the introduction of those circuits.

### 1.2 Study two - Modification of enclosures for single-housed monkeys

Although social housing is the rule, on occasions there are barriers to its implementation. Sometimes, for reasons of behavioural incompatibility or on veterinary advice, it is necessary to house animals in individual cages. While animals are alone in their cage, they are in the same room as other monkeys and have visual, olfactory and auditory access to their neighbours. Single-housed monkeys are known to be less active and have more stereotypical behaviours than monkeys housed in groups (26,27). However, additional enrichment is commonly provided to these animals to alleviate these adverse effects (28). Inspired by the work of Disarbois and Duhamel on cynomolgus monkeys (15), we tested whether providing more visual contact between rhesus monkeys in adjacent enclosures by replacing opaque panels with transparent ones could further improve animals’ welfare. We hypothesize that offering them the possibility of having more direct visual interactions with their conspecific could help reduce stereotypical behaviours like pacing, and promote social interactions through a form of simulated grooming.

We implemented this addition of transparent panels in two enclosures. The first one was placed in between enclosures of two single-housed animals; the second was placed in between cages of a single-housed animal and of pair-housed animals. We focused our analysis on the behaviour of the 3 single-housed monkeys in 23 and 39 sessions of behavioural observations for monkey Qu and for monkeys Dai and Ra respectively. First in typical setting conditions, with the opaque panel (condition *Opaque*), then during the introduction of the transparent panel (a couple of hours a day for several weeks) (condition *Habituation*) and finally when the transparent panel was set permanently (condition *Transparent*). Data from only two out of three animals were collected in the condition *Transparent*, as one animal was moved to a different room during the project.

For all monkeys, we observed a small increase of aggressive behaviour at the time of the introduction of the transparent panel that rapidly decreased over time (Fig. 3A). More importantly, we observed across sessions a decrease of the time spent pacing after the introduction of the transparent panel (Fig. 3A). The overall pacing time was significantly lower during the habituation phase compared to the condition *Opaque* (p-value = 0.0006, with Bonferroni correction) (Fig. 3B). This benefit was long lasting as it sustains after the permanent addition of the transparent panel (condition *Transparent*).

**Figure 3:**
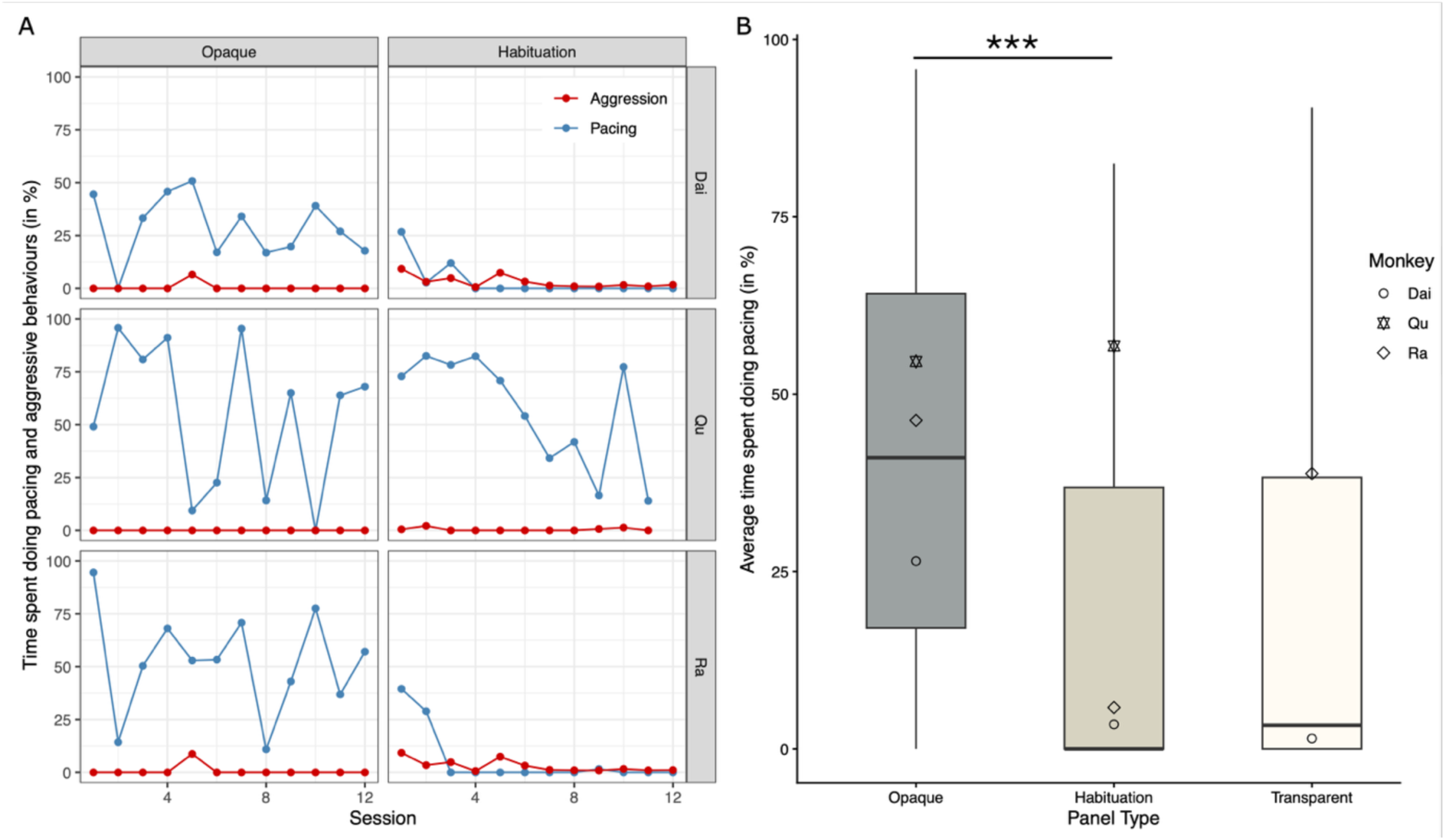
Effect of the panel type on the time spent having abnormal and aggressive behaviours across 3 monkeys. A. Time of pacing and aggressive behaviours across 10 minutes sessions during the typical condition with the opaque panel (condition Opaque) (left column) and condition Habituation with the transparent panel (condition Habituation) (right column). Pacing is represented in blue. Aggressive behaviour is represented in red. B. Average time pacing of three monkeys during three panel conditions (opaque, habituation, transparent). In comparison to the condition Opaque, it shows a significant decrease during the condition Habituation (p-value = 0.0006), maintained during the condition Transparent (p-value = 0.4520, habituation compared to condition Transparent).

In addition, we observed that over time recurrent virtual grooming sessions occurred in between the two single-housed animals (See Supplementary Fig.1). Those two animals were recently successfully paired following the procedure. This pseudo-grooming behaviour was not observed between monkey Qu and its neighbours.

By implementing these two simple structural enrichments, we managed to influence the expression of natural behaviour in our socially housed and single housed monkeys by increasing the number of appeasing non-social behaviours and decreasing the duration of abnormal behaviours respectively.

## 2. Enrichment - Relevance and impact of manipulable objects

### 2.1 Study three - Impact of enrichments on socially and single housed monkeys’ time budget

Our enrichment programme includes the daily provision of manipulable objects to monkeys. Objects are added from one day to the next, and only removed from the enclosures once a week. In a first step to optimize this programme, we tested its efficacy by measuring the changes on the monkeys’ time budget associated with 3 different versions of the programme. We compared three conditions: a week without the distribution of any additional objects (condition *Without*), a week with the distribution of known objects (condition *Known*) every day and a week with the distribution of unknown objects (condition *New*) every day. We conducted behavioural observations 3 times a day over consecutive days in a group of 3 monkeys and in 2 single-housed animals. More specifically, we measured the time they spent engaged in ten different behavioural categories: abnormal behaviour, affiliative and neutral social behaviour, aggressive social behaviour, feeding behaviour, interacting with cage elements, moving around, resting, self-grooming/scratching, using objects and other behaviours (Fig. 4).

**Figure 4:**
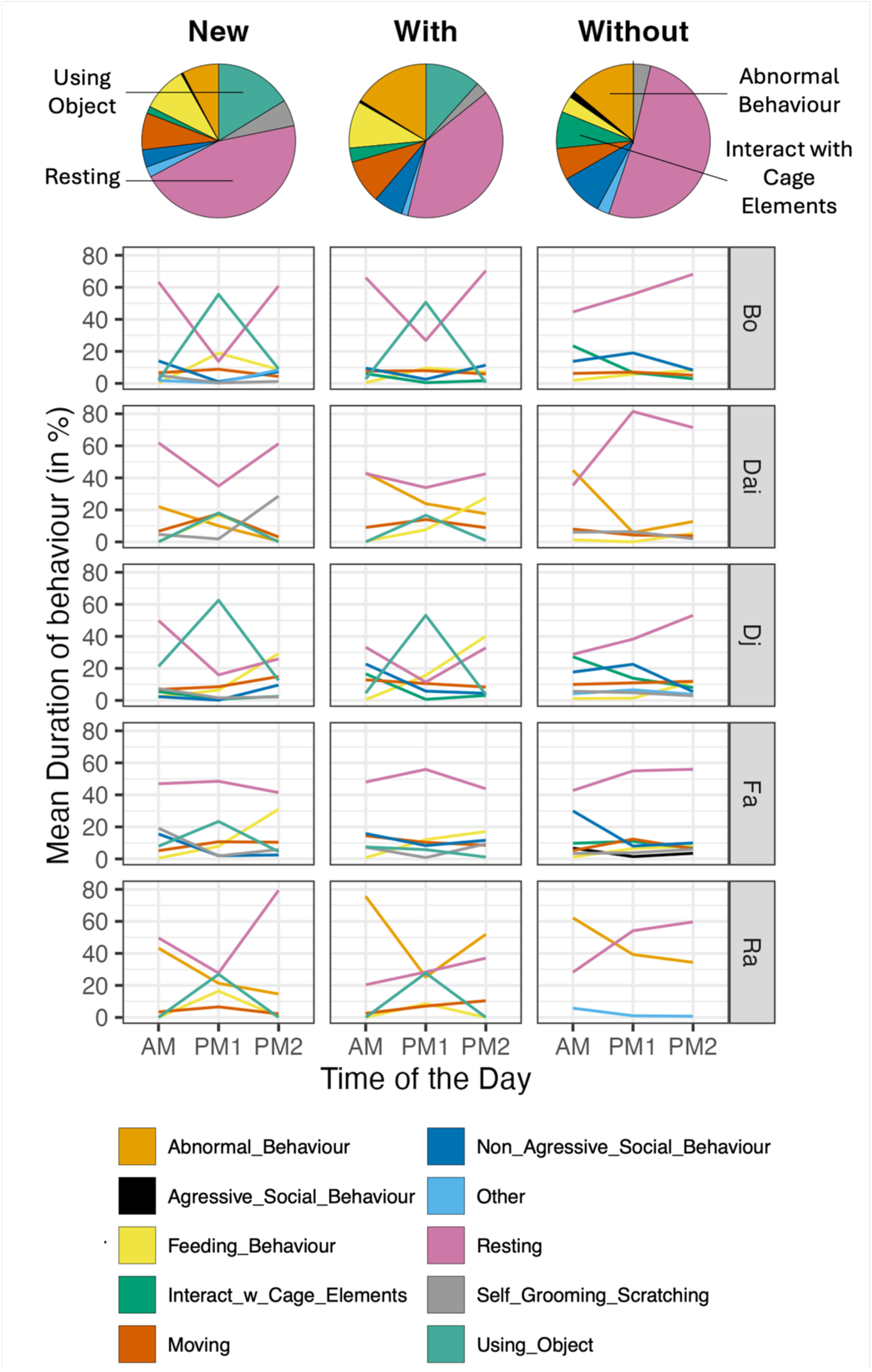
Evaluation of the daily time budgets of a group of three males and two single-housed monkeys, tested in three different environment conditions, enriched with unknown objects (New), enriched with known objects (Known) and not enriched (Without). The behaviours represented are shown only if they occur more than 5% of the time in at list one of the three time points we measured (AM, PM1, PM2). This figure shows different patterns influenced by individuals, condition and time within the day.

At the group level, PERMANOVA revealed significant effects of the time of the day and of the experimental condition (p-values < 0.0001). Timewise, the 3 animals in a social group were less active in the morning, prior to the distribution of objects, compared with early or late afternoon. The 2 single-housed animals displayed more stereotypical behaviours in the morning than during the rest of the day. We then conducted pairwise post-hoc comparisons by testing each pair of experimental conditions separately with Bonferroni correction. The three environment conditions (*New*, *Known*, *Without*) showed significantly different behavioural profiles from one another (pairwise permanova; *New* vs *Known* F(1,145) = 2.893, p-value = 0.008; *New* vs *Without* F(1,140) = 14.874, p-value < 0.001; *Known* vs *Without* F(1,145) = 9.457, p-value < 0.001). However, these results should be interpreted with caution. Ordination (PCoA) showed substantial overlap between groups, indicating that effect sizes were small (R² = 0.077 and R² = 0.114, respectively). Additional Betadisper tests indicated heterogeneous within-group dispersions. Altogether these two additional analyses are pointing to the large inter-individual variability in our data.

To better understand this interindividual variability, we then conducted an analysis at the subject level (Fig. 5; Supplementary Table 1; Supplementary Fig. 2). We observed notably that all individuals used the manipulable objects we provided but only one individual (Fa) used more the new objects than the known objects (Fig. 5; Supplementary Table 1; Supplementary Fig. 2). Although single-housed animals did not use new objects more than known ones, they were sensitive to novelty as the introduction of new objects is associated with less abnormal behaviours (Fig. 5).

**Figure 5:**
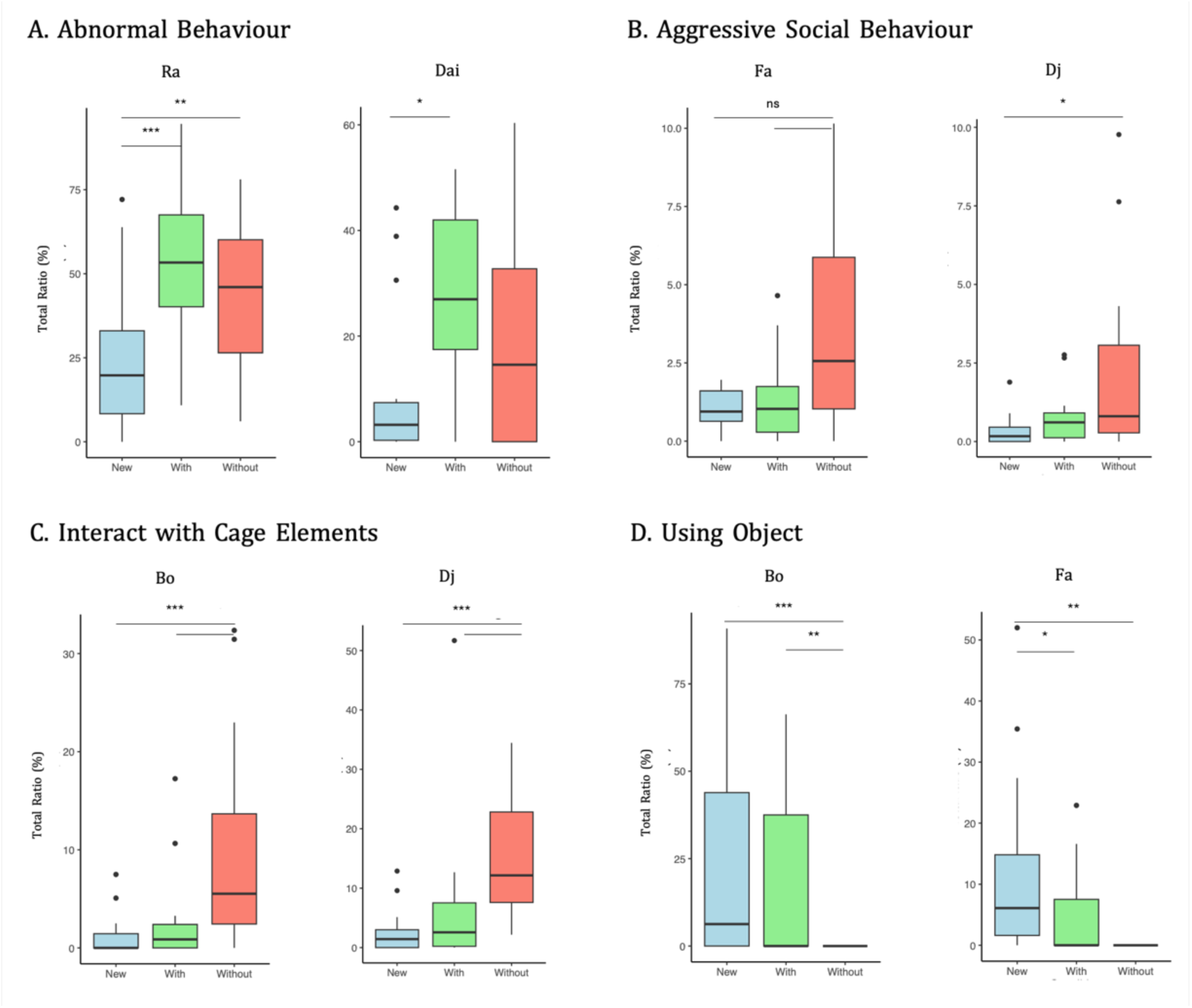
Significant behaviours for all five monkeys. A. Abnormal Behaviour i) Monkey Ra: more in conditions Known and Without compared to New (p-value = 0.0002 and p-value = 0.0057 respectively) ii) Monkey Da: more in condition Known compared to New (p-value = 0.0161). B. Aggressive Social Behaviour i) Monkey Fa: Tendency but not significantly more in Without compared to Known and New (p-value = 0.1011 and p-value = 0.1014 respectively) ii) Monkey Dj: more in condition Without compared to New (p-value = 0.026). C. Interact with Cage Elements i) Monkey Bo: more in condition Without compared to Known and New (p-value = 0.0005 and p-value = 0.0002 respectively) ii) Monkey Dj: more in condition Without compared to Known and New (p-value < 0.0001 and p-value < 0.0001 respectively). D. Using Object i) Monkey Bo: more in condition Known and New compared to Without (p-value = 0.0014 and p-value = 0.0003 respectively), no difference between Known and New (p-value = 0.7015) ii) Monkey Fa: more in condition New compared to Without and Known (p-value = 0.0012 and p-value = 0.0480 respectively), no difference between Known and Without (p-value = 0.3757). Results from monkeys Dj, Ra and Dai are similar to Bo and shown on supplementary figure 2.

When objects were not available, two animals in the social group (Bo, Dj) interacted more with the structural enrichment left available in the enclosures (e.g. swing) (Fig. 5). In addition, we observed an increase of aggressive behaviour in two animals of the social group when manipulable objects were absent, with one animal displaying more aggressivity (Dj), while another individual was more targeted (Fa) (Fig. 5). We did not observe changes in affiliative and neutral social behaviour in none of the three conditions.

Finally, we did observe that the subordinate animal (Fa) had less access to enrichment than dominant macaques Bo and Djo. The usage of the objects we distribute daily appeared to be delayed. Due to our sample, this phenomenon was difficult to capture in this analysis.

### 2.2 Study four - Object relevance for individualised enrichment programmes

In a second step to optimize our enrichment programme, we aimed to identify potential preferences for specific manipulable objects at the individual level.

We presented twenty rhesus monkeys with four different objects (silver ball, kong, bottle, puzzle or silver ball, fire hose, noise ball, tube depending on the research colonies) and we measured how long they interacted with each item during a 10-minutes observation session. Once again, these results showed large inter individual differences (Fig 6).

**Figure 6:**
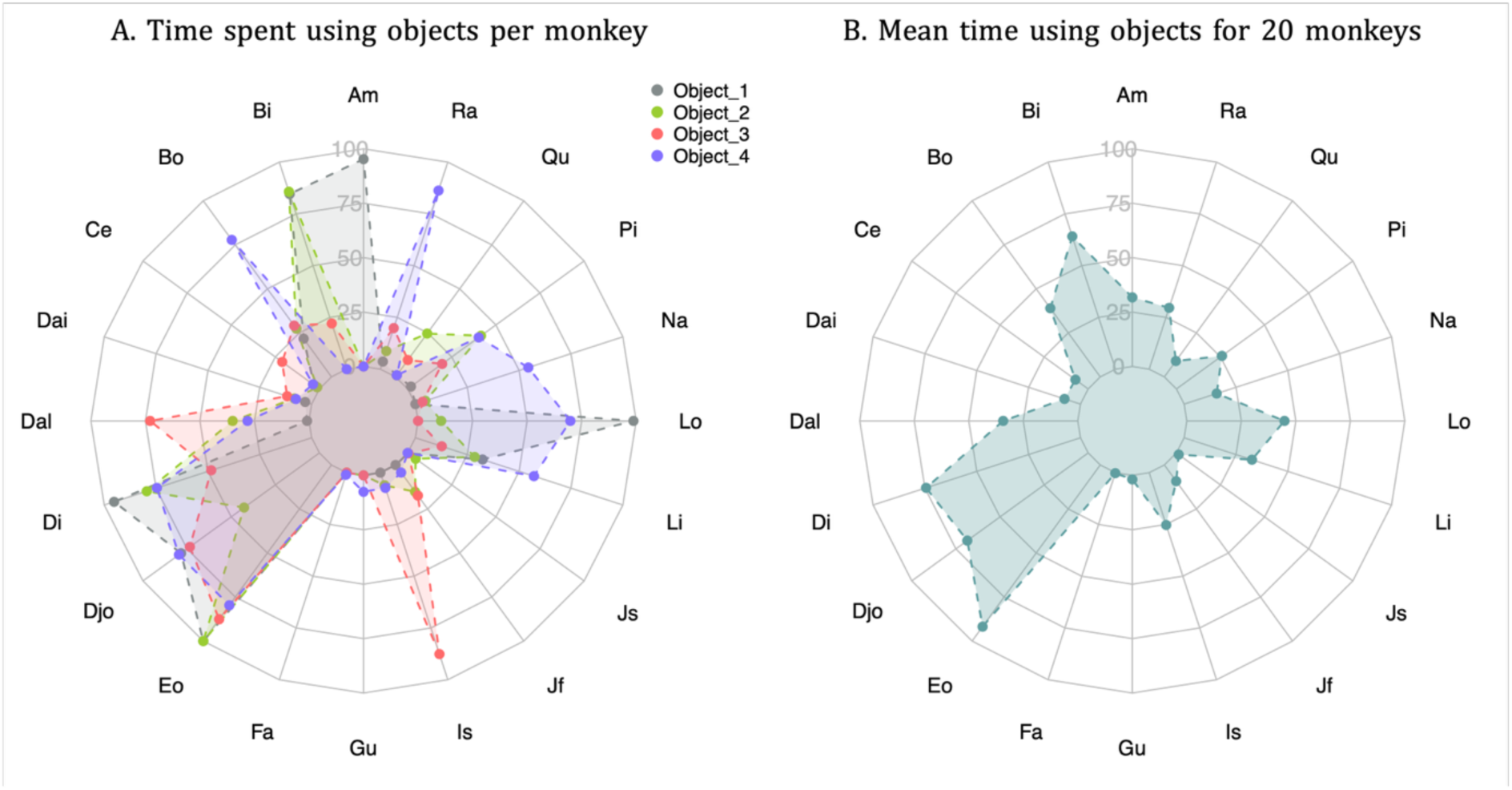
A. Representation of the time spent using 4 objects for each monkeys (n = 20) in percent showing important interindividual preferences and variability (Obj4 missing for Am and Bi). Objects tested were different for the three colonies: Colonies 1 and 2: Object_1 = silver ball, Object_2 = kong, Object_3 = bottle, Object_4 = puzzle; Colony 3: Object_1 = silver ball, Object_2 = fire hose, Object_3 = noise ball, Object_4 = tube. B. Mean time using objects during 10 minutes observation for 20 monkeys. Three or four objects were tested for each individual. This figure shows a strong interindividual variability.

We then checked whether sociobiological variables (sex, age, social status, group size) could explain this interindividual variability. We used two generalized linear mixed models with Beta Zero-One-Inflated distribution to test the potential effect of age, sex, and the size of the social group. None of the predictors reached the statistical significance (age: p-value = 0.40; sex: p-value = 0.613; social group size: p-value = 0.452). For the socially housed animals, we ran an additional analysis including social status. None of the predictors reached statistical significance (age: p-value = 0.816; sex: p-value = 0.808; social group size: p-value = 0.904; social status: p-value = 0.933). Note that only 13 subjects were included in this second analysis, reducing the statistical power of our analysis.

## Discussion

In this project, we assessed effects of changes to monkey enclosures and of the effects of our enrichment programme on animals’ behaviour. Firstly, we showed that by adding new paths, or circuits, to animal enclosures, we were able to change dynamics in the social groups we tested. Although we did not impact appeasing or aggressive social behaviours, we reduced the occurrence of injuries requiring veterinary intervention and increased the expression of appeasing non-social behaviours. Adding circuits is not preventing social tension between macaques but it impacts on the way animals deal with it. For submissive animals, it offers several options to escape aggressive dominant animals. Secondly, we removed some visual barriers between enclosures. Visual barriers are traditionally implemented to reduce social tension between neighbouring groups (29). Here we tested whether transparent panels between neighbouring enclosures of single-housed monkeys and between single-housed and pair-housed animals could have a beneficial effect. While we obtained positive effects of the introduction of the transparent panel in the long term, it caused initially a short-lived increase of aggressivity between the animals we tested. Over the long term, our results showed that the use of the transparent panels was associated with a reduction of pacing in single-housed animals. One could interpret the benefit of a transparent barrier as providing visual information that might help resolve sources of uncertainty coming from the noise in the adjacent enclosure that cannot be stopped. Furthermore, we also observed that a transparent barrier could also lead to virtual social interactions between the two neighbouring single-housed animals. This phenomenon had been previously observed in cynomolgus monkeys (15). In addition to reducing sensory uncertainty, a transparent barrier could activate the peripersonal space system, a set of brain regions that contain neurons with visuo-tactile properties and that respond to objects and agents positioned close to the body where interactions are facilitated (30–32). Neurons of this system do respond to body proximity even in the presence of a transparent barrier (33,34). Interestingly, we were then able to successfully pair the two animals that displayed virtual interactions through the transparent panel. A follow-up study could determine whether the emergence of such behaviour is a predictor of the success of a pairing between animals.

In addition, we evaluated the benefit of our enrichment programme. Our results confirmed the benefits of the distribution of manipulable objects on animals’ behaviours in both single-housed and socially-housed monkeys. Their presence is associated with more active animals in social-housed monkeys, while their absence is associated with an increase in aggression. Importantly, our results showed benefits of new objects on single-housed monkeys with a reduction of stereotypical behaviours. As the interest for a given object could decrease over time, it would be important to renew manipulable objects on a regular basis to ensure that enrichment maintains its efficacy. Future study could compare the value of a cumulative enrichment programme with a non-cumulative programme in which enrichment is changed on a daily basis.

Our results also revealed the huge inter-individual variability in response to manipulable objects. Previous studies have already shown that animal reactions will vary depending on the species or the social environment (35–37). In our study, this variability could not be explained by age, sex, or social factors such as social status or social group size. Instead it appears that how an animal will react to a manipulable object is a reflection of the personality traits of each individual (37). Therefore, objects used in enrichment programmes should be evaluated on a subject-by-subject basis to ensure appropriate selection of objects in any enrichment programme. While animals lose interest in objects in within hours following their distribution, and that little reuse of objects is generally observed (38,39), some of our observations suggest that a cumulative protocol gives the opportunity to subordinate monkeys to access objects long after their distribution and once dominant animals have finished using them. Future study will aim at specifically addressing this question on a larger sample. These results show also how important it is to regularly change the objects we provide monkeys with and question the frequency of distribution of each object. Finally, providing a sufficient number of objects spread across the environment will allow for better access for subordinate monkeys and limit the possibility of aggressions (40,41).

Overall, our data support the development of evidence-based enrichment programmes. The limited number of animals speaks for the need of a cooperative approach to this question that should be tackled at the level of national or international consortiums such as Biosimia, Eusimia or Simco.

## Ethics / Funding

This project was discussed with the Animal Welfare Body of the laboratories in which animals were housed. The data presented in this article were collected in the context of ethically approved protocols by French Animal Experimentation Ethics Committee #42 (CELYNE) that followed guidelines of the European Community on animal care (European Community Council Directive 2010/63/EU).

This work was supported by the Agence Nationale de la Recherche (ANR-22-CE37-0021; ANR-21-CE37-0016; ANR-18-CE37-0016) and the Réseau médical & scientifique de l’Institut de Neuromodulation, INM, GHU Paris Psychiatrie et Neurosciences. VML was supported by Ministère de l’Éducation nationale, de la Recherche et de la Technologie (MENRT) and Fondation pour la Recherche Médicale PhD studentships (FRM: FDT202504020240).

## Acknowledgments

We are very grateful for the care afforded to the animals by the veterinary, caregiving and technical staff. We thank J Lachaud, L Delabuxière, J Lavignas, M Filiptchenko, M Vivet, O Jourjon, A Chekki and E Vigier for their help with looking after the animals.

## Supplementary

**Supplementary Figure 1:**
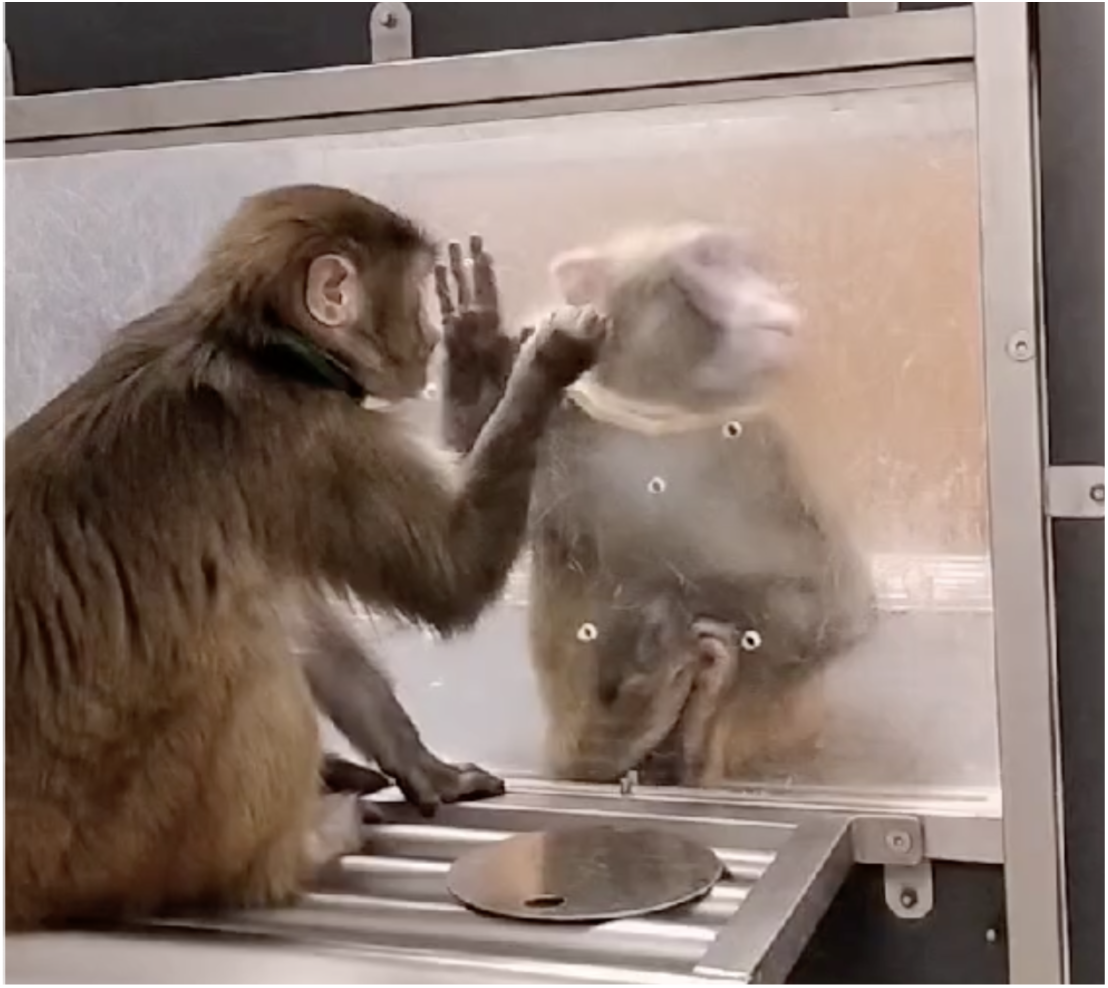
Photo of monkey Dai displaying virtual grooming towards monkey Ra through the transparent panel.

**Supplementary Figure 2:**
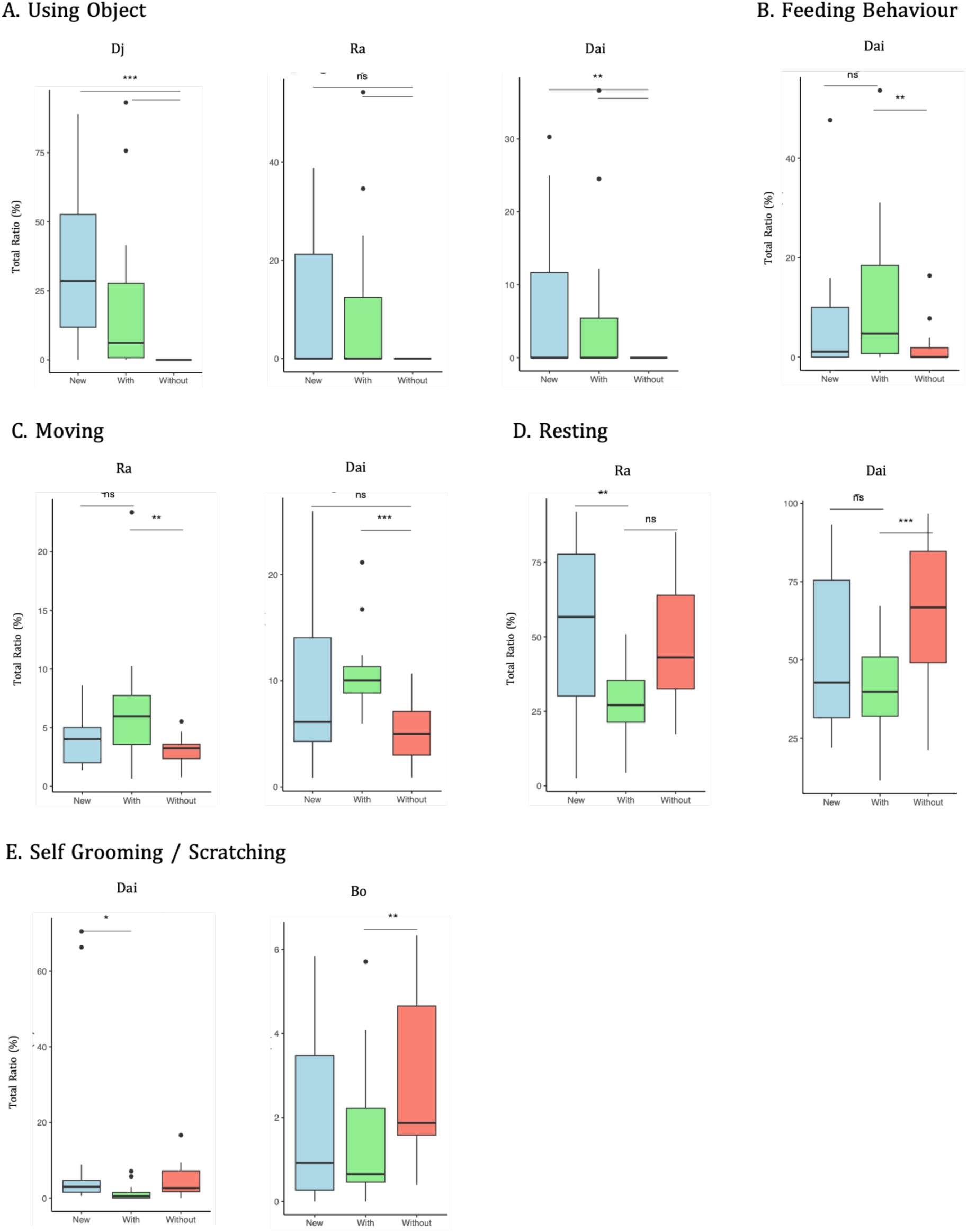
Significant behaviours for all five monkeys. A. Using object i) Monkey Dj: more in condition Known and New compared to Without (p-value = 0.0006 and p-value = 0.0010 respectively), no difference between Known and New (p-value = 0.4940) ii) Monkey Ra: Tendency but not significantly more in conditions Known and New compared to Without (p-value = 0.0750 and p-value = 0.0794 respectively) but model didn’t pass all validation criteria iii) Monkey Dai: more in condition Known and New compared to Without (p-value = 0.0041 and p-value = 0.0045 respectively), no difference between Known and New (p-value = 0.9996) but the model didn’t pass all validation criteria. Note that monkeys Bo and Fa are shown in the main text figure. B. Feeding behaviour, Monkey Dai: more in condition Known compared to Without (p-value = 0.0069) and tendency but not significantly more in condition Known compared to New (p-value = 0.0749). C. Moving i) Monkey Ra: more in condition Known compared to Without (p-value = 0.0021) ii) Monkey Dai: more in condition Known compared to Without (p-value < 0.0001) and tendency but not significantly more in condition New compared to Without (p-value = 0.0670). D. Resting i) Monkey Ra: more in condition New compared to Known (p-value = 0.0016) ii) Monkey Dai: more in condition Without compared to Known (p-value = 0.0004). E. Self-grooming / scratching i) Monkey Dai: more in condition New compared to Known (p-value = 0.0378) but model didn’t pass all validation criteria ii) Monkey Bo: more in condition Without compared to Known (p-value = 0.0045).

---

**Supplementary Table 1:** Selected models for each behaviour and each monkey, based on AIC and quality checks.

| ID Monkey | Behaviour | Selected Model |
| --- | --- | --- |
| Bo | Abnormal behaviour | beta_additif |
| Bo | Affiliative and neutral social behaviour | beta_additif |
| Bo | Aggressive social behaviour | beta_additif |
| Bo | Feeding behaviour | beta_additif |
| Bo | Interacting with cage elements | beta_additif |
| Bo | Moving around | beta_additif |
| Bo | Resting | beta_additif |
| Bo | Self-grooming/scratching | beta_interaction |
| Bo | Using object | beta_additif |
| Bo | Other | beta_additif |
| Fa | Abnormal behaviour | beta_additif |
| Fa | Affiliative and neutral social behaviour | beta_additif |
| Fa | Aggressive social behaviour | beta_interaction |
| Fa | Feeding behaviour | beta_additif |
| Fa | Interacting with cage elements | beta_additif |
| Fa | Moving around | beta_interaction |
| Fa | Resting | beta_additif |
| Fa | Self-grooming/scratching | beta_additif |
| Fa | Using object | zi_beta_additif |
| Fa | Other | beta_additif |
| Dj | Abnormal behaviour | beta_interaction |
| Dj | Affiliative and neutral social behaviour | beta_additif |
| Dj | Aggressive social behaviour | beta_additif |
| Dj | Feeding behaviour | beta_additif |
| Dj | Interacting with cage elements | beta_additif |
| Dj | Moving around | beta_additif |
| Dj | Resting | beta_interaction |
| Dj | Self-grooming/scratching | beta_additif |
| Dj | Using object | beta_interaction |
| Dj | Other | beta_additif |
| Ra | Abnormal behaviour | beta_additif |
| Ra | Affiliative and neutral social behaviour | beta_interaction |
| Ra | Aggressive social behaviour | beta_additif |
| Ra | Feeding behaviour | beta_additif |
| Ra | Interacting with cage elements | beta_additif |
| Ra | Moving around | beta_additif |
| Ra | Resting | beta_interaction |
| Ra | Self-grooming/scratching | beta_additif |
| Ra | Using object | beta_interaction |
| Ra | Other | beta_additif |
| Dai | Abnormal behaviour | beta_additif |
| Dai | Affiliative and neutral social behaviour | beta_additif |
| Dai | Aggressive social behaviour | beta_additif |
| Dai | Feeding behaviour | beta_additif |
| Dai | Interacting with cage elements | beta_additif |
| Dai | Moving around | beta_interaction |
| Dai | Resting | beta_interaction |
| Dai | Self-grooming/scratching | gaussian_sqrt |
| Dai | Using object | beta_interaction |
| Dai | Other | beta_additif |

## References

1. Roelfsema PR, Treue S. Basic neuroscience research with nonhuman primates: a small but indispensable component of biomedical research. Neuron. 2014 Jun 18;82(6):1200–4.

2. Procyk E, Ancé P, Badin RA, Balansard I, Boraud T, Brochier T, et al. Les primates non-humains et la recherche biomédicale en France. hal-03755028 [Internet]. 2020 [cited 2026 May 5]; Available from: https://hal.science/hal-03755028v1

3. Phillips KA, Bales KL, Capitanio JP, Conley A, Czoty PW,’t Hart BA, et al. Why primate models matter. Am J Primatol. 2014 Sep;76(9):801–27.

4. Homberg JR, Adan RAH, Alenina N, Asiminas A, Bader M, Beckers T, et al. The continued need for animals to advance brain research. Neuron. 2021 Aug 4;109(15):2374–9.

5. Feng G, Jensen FE, Greely HT, Okano H, Treue S, Roberts AC, et al. Opportunities and limitations of genetically modified nonhuman primate models for neuroscience research. Proc Natl Acad Sci USA. 2020 Sep 29;117(39):24022–31.

6. Morel-Latour V, Ravel S, Dirheimer M, Sallet J. Aspects éthiques et cadres juridiques de la recherche en neurosciences comportementales chez les primates non-humains. Primatologie. 2025;16.

7. Lutz CK, Novak MA. Environmental enrichment for nonhuman primates: theory and application. ILAR J. 2005;46(2):178–91.

8. Morel-Latour V, Ravel S, Dirheimer M, Sallet J. Bientraitance et bien-être des primates non-humains en recherche en neurosciences comportementales: mise en œuvre du principe de raffinement. Primatologie. 2025;16.

9. Buchanan-Smith HM. Environmental enrichment for primates in laboratories. Adv Sci Res. 2011 Jun 14;5(1):41–56.

10. Bloomstrand M, Riddle K, Alford P, Maple T.L. Objective Evaluation of a Behavioral Enrichment Device for Captive Chimpanzees (Pan troglodytes). Zoo Biol. 1986;5:293–300.

11. Kessel AL, Brent L. Cage toys reduce abnormal behavior in individually housed pigtail macaques. J Appl Anim Welf Sci. 1998;1(3):227–34.

12. Akhund-Zade J, Ho S, O’Leary C, de Bivort B. The effect of environmental enrichment on behavioral variability depends on genotype, behavior, and type of enrichment. J Exp Biol. 2019 Oct 8;222(Pt 19).

13. Miller LJ, Vicino GA, Sheftel J, Lauderdale LK. Behavioral diversity as a potential indicator of positive animal welfare. Animals (Basel). 2020 Jul 16;10(7).

14. Schapiro SJ, Bloomsmith MA, Kessel AL, Shively CA. Effects of enrichment and housing on cortisol response in juvenile rhesus monkeys. Appl Anim Behav Sci. 1993 Aug;37(3):251–63.

15. Disarbois E, Duhamel J-R. Virtual social grooming in macaques and its psychophysiological effects. Sci Rep. 2024 May 22;14(1):11697.

16. Albanese V, Kuan M, Accorsi PA, Berardi R, Marliani G. Evaluation of an enrichment programme for a colony of long-tailed macaques (Macaca fascicularis) in a rescue centre. Primates. 2021 Jul;62(4):585–93.

17. Márquez-Arias A, Santillán-Doherty AM, Arenas-Rosas RV, Gasca-Matías MP, Muñoz-Delgado J. Environmental enrichment for captive stumptail macaques (Macaca arctoides). J Med Primatol. 2010 Feb;39(1):32–40.

18. Bryant CE, Rupniak NM, Iversen SD. Effects of different environmental enrichment devices on cage stereotypies and autoaggression in captive cynomolgus monkeys. J Med Primatol. 1988;17(5):257–69.

19. de Waal FBM. The myth of a simple relation between space and aggression in captive primates. Zoo Biol. 1989;8(S1):141–8.

20. Izzo GN, Bashaw MJ, Campbell JB. Enrichment and individual differences affect welfare indicators in squirrel monkeys (Saimiri sciureus). J Comp Psychol. 2011 Aug;125(3):347–52.

21. Sallet J, Mars RB, Noonan MP, Andersson JL, O’Reilly JX, Jbabdi S, et al. Social network size affects neural circuits in macaques. Science. 2011 Nov 4;334(6056):697–700.

22. Sallet J, Emberton A, Wood J, Rushworth M. Impact of internal and external factors on prosocial choices in rhesus macaques. Philos Trans R Soc Lond B Biol Sci. 2021 Mar;376(1819):20190678.

23. Noonan MP, Sallet J, Mars RB, Neubert FX, O’Reilly JX, Andersson JL, et al. A neural circuit covarying with social hierarchy in macaques. PLoS Biol. 2014 Sep 2;12(9):e1001940.

24. Fedigan LM. Primates Paradigms: sex roles and social bonds. University of Chicago Press edition 1992. The University of Chicago Press, Chicago, USA; 1992.

25. Friard O, Gamba M. BORIS: a free, versatile open-source event-logging software for video/audio coding and live observations. Methods Ecol Evol. 2016 Nov;7(11):1325–30.

26. Schapiro SJ, Bloomsmith MA, Porter LM, Suarez SA. Enrichment effects on rhesus monkeys successively housed singly, in pairs, and in groups. Appl Anim Behav Sci. 1996 Jul;48(3–4):159–71.

27. Bellanca RU, Crockett CM. Factors predicting increased incidence of abnormal behavior in male pigtailed macaques. Am J Primatol. 2002 Oct;58(2):57–69.

28. Griffis CM, Martin AL, Perlman JE, Bloomsmith MA. Play caging benefits the behavior of singly housed laboratory rhesus macaques (Macaca mulatta). J Am Assoc Lab Anim Sci. 2013 Sep;52(5):534–40.

29. Reinhardt V. Common husbandry-related variables in biomedical research with animals. Lab Anim. 2004 Jul;38(3):213–35.

30. Livi A, Lanzilotto M, Maranesi M, Fogassi L, Rizzolatti G, Bonini L. Agent-based representations of objects and actions in the monkey pre-supplementary motor area. Proc Natl Acad Sci USA. 2019 Feb 12;116(7):2691–700.

31. Brozzoli C, Makin TR, Cardinali L, Holmes NP, Farnè A. Peripersonal Space: A Multisensory Interface for Body–Object Interactions. In: Murray MM, Wallace MT, editors. The Neural Bases of Multisensory Processes. Boca Raton (FL): CRC Press/Taylor & Francis; 2012.

32. Bogdanova OV, Bogdanov VB, Dureux A, Farnè A, Hadj-Bouziane F. The Peripersonal Space in a social world. Cortex. 2021 Sep;142:28–46.

33. Caggiano V, Fogassi L, Rizzolatti G, Thier P, Casile A. Mirror neurons differentially encode the peripersonal and extrapersonal space of monkeys. Science. 2009 Apr 17;324(5925):403–6.

34. Farnè A, Demattè ML, Làdavas E. Beyond the window: multisensory representation of peripersonal space across a transparent barrier. Int J Psychophysiol. 2003 Oct;50(1–2):51–61.

35. Addessi E, Chiarotti F, Visalberghi E, Anzenberger G. Response to novel food and the role of social influences in common marmosets (Callithrix jacchus) and Goeldi’s monkeys (Callimico goeldii). Am J Primatol. 2007 Nov;69(11):1210–22.

36. Englerova K, Klement D, Frynta D, Rokyta R, Nekovarova T. Reactions to novel objects in monkeys: what does it mean to be neophobic? Primates. 2019 Jul;60(4):347–53.

37. Šlipogor V, Graf C, Massen JJM, Bugnyar T. Personality and social environment predict cognitive performance in common marmosets (Callithrix jacchus). Sci Rep. 2022 May 5;12(1):6702.

38. Line SW, Morgan KN. The effects of two novel objects on the behavior of singly caged adult rhesus macaques. Lab Anim Sci. 1991 Aug;41(4):365–9.

39. Line SW, Morgan KN, Markowitz H. Simple toys do not alter the behavior of aged rhesus monkeys. Zoo Biol. 1991 Jan;10(6):473–84.

40. Newberry RC. Environmental enrichment: Increasing the biological relevance of captive environments. Appl Anim Behav Sci. 1995 Sep;44(2–4):229–43.

41. Honess PE, Marin CM. Enrichment and aggression in primates. Neurosci Biobehav Rev. 2006;30(3):413–36.

